# Epitope Prediction of Antigen Protein using Attention-Based LSTM Network

**DOI:** 10.1101/2020.07.27.224121

**Authors:** Toshiaki Noumi, Seiichi Inoue, Haruka Fujita, Kugatsu Sadamitsu, Makoto Sakaguchi, Akiko Tenma, Hironori Nakagami

## Abstract

B-cells inducing antigen-specific immune responses in vivo produce large amounts of antigen-specific antibodies by recognizing the subregions (epitope regions) of antigen proteins. They can inhibit their functioning by binding antibodies to antigen proteins. Predicting of epitope regions is beneficial for the design and development of vaccines aimed to induce antigen-specific antibody production. However, prediction accuracy requires improvement. The conventional epitope region prediction methods have focused only on the target sequence in the amino acid sequences of an entire antigen protein and have not thoroughly considered its sequence and features as a whole. In the present paper, we propose a deep learning method based on short-term memory with an attention mechanism to consider the characteristics of a whole antigen protein in addition to the target sequence. The proposed method achieves better accuracy compared with the conventional method in the experimental prediction of epitope regions using the data from the immune epitope database.

## 1. Introduction

B-cells, which induce antigen-specific immune responses in vivo, produce large amounts of antigen-specific antibodies by recognizing the subregions (epitope regions) of antigen proteins. Antibodies can inhibit the functioning of an antigen protein by binding it to its epitope region [1]. A substance that mimics the structure and function of an epitope can be considered as a “vaccine” to an organism that is aimed to induce specific antibodies in vivo. Various studies focused on epitopes have been conducted to design safe and effective vaccines. Conducting the three-dimensional structural analysis of antibody-antigen complexes by X-ray [2] or nuclear magnetic reasonance (NMR) spectroscopy [3] is deemed as the most reliable way to identify epitopes recognized by B-cells. However, this procedure is expensive in terms of time, cost, and labor. Therefore, to address this problem, the computer-based epitope prediction has been introduced. Recently, various machine learning methods have been proposed [4], [5]. Although their performance has been improved, the employed features are limited to those associated with the target amino acid sequence, and therefore, the representation capability of such models is insufficient.

In the present paper, we propose a method for B-cell epitope prediction using an attention-based long short-term memory network (LSTM) [6] (a deep learning approach) to incorporate not only the target amino acid sequence but also the long-range features of a whole protein. In addition, the attention mechanism [7] is realized to enable automatically estimating the points to be emphasized while predicting each amino acid in and out of candidate epitopes. Furthermore, to address the problem of data sparseness, we extend the method to enable the simultaneous consideration of the structural and chemical features of an entire antigen protein in a deep learning network.

We demonstrate that the proposed method achieves better prediction accuracy compared with the existing method BepiPred-2.0 [4] through implementing it in the experiments on immune epitope database (IEDB) that is a public database of immune epitopes [8], [9]. Dataset and some basic programs used in the present study will be made available to the public ^*1^.

## 2. Task definition

### 2.1 Task definition

In the present study, the prediction of B-cell epitopes was based on the long-chain amino acid sequences that constituted antigenic proteins. The epitope regions registered in IEDB approximately ranged from 5 to 20 amino acids [8], [9], [10]. Although the proposed method could be applied regardless of the length of an epitope region, in the problem setting of this research, we limited the length of a candidate epitope peptide (corresponding to the short amino acid sequences) to 8-14 amino acids. We addressed the problem of classifying of these peptides into two categories: with antibody inducing activity (positive) and without such activity (negative).

### 2.2 Employed dataset

Information on whether or not an amino acid peptide exhibited antibody-inducing activity (marked by an activity label) could be obtained from IEDB [8], [9], [11], which was used in many previous studies. Accordingly, this information was used as the label data. We also obtained the epitope candidate amino acid sequences (peptides) and the activity label data from the B-cell epitope data provided in IEDB. The presented antibody proteins were restricted to IgG that constituted the most recorded type in IEDB. For convenience, we excluded records representing different quantitative measures of antibody activity for the same peptide from experiments. The epitope data obtained from IEDB corresponded to the five types of activity: “Positive-High,” “Positive-Intermediate,” “Positive-Low,” “Positive,” and “Negative.” However, due to the limited number of data elements marked with the “Positive-High,” “Positive-Intermediate,” and “Positive-Low” labels, we equally considered these labels as “Positive”, thereby attributing the task to a binary estimation. In Table 1, we represented “the number of peptides,” “the number of proteins,” and “the ratio of positives” in the considered data including each length. Notably, the number of significant digits was three. As shown in culumn “the ratio of poitives” in Table 1, we noted that there was a difference between the positive ratio values corresponding to various lengths.

**Table 1.** Statistics of the used dataset.

| length | Number of peptides | Number of proteins | Ratio of positives |
| --- | --- | --- | --- |
| 8 | 2271 | 135 | 0.197 |
| 9 | 242 | 128 | 0.675 |
| 10 | 3276 | 180 | 0.227 |
| 11 | 346 | 125 | 0.204 |
| 12 | 758 | 172 | 0.492 |
| 13 | 245 | 186 | 0.736 |
| 14 | 337 | 127 | 0.458 |
| total | 7475 | 646 | 0.293 |

Concerning this population, we extracted the dataset corresponding to 10 different random states with the ratio of the learning data to the evaluation data at approximately 10:1. The data were split without duplication across all sets. In each dataset, there were no duplications of proteins across train and test data. Furthermore, we excluded the peptides with high homology from each dataset, using CD-HIT [13]. CD-HIT is the tool, which determines degree of homology between two sequences. In this study, we calculated degree of homology for all combinations of peptide sequences in the train and test dataset by CD-HIT. From previous study [14], we set 40% as a maximum degree of homology and used the data as no homology sequences data.

## 3. Related work and issues

The early computer-based B-cell epitope prediction methods focused solely on the physicochemical properties of the amino acids constituting protein [12]. In contrast to these manually derived predictions based on specific indices, machine learning methods incorporating the information from the amino acid sequences themselves achieved high performance [4], [5]. Various methods based on machine learning algorithms were proposed, including those using random Forest (published tool BepiPred-2.0 [4]), support vector machine (SVM) and k-neighborhood method (published tool LBtope) [5]. Although the performance of these methods could not be unconditionally compared due to different dataset used in the experiments, BepiPred-2.0 achieved the best results [4] in that there is the significant difference in AUC between BepiPred-2.0 and others. In several previous studies, short amino acid sequences (1-3 amino acids), before and after the target peptide, were added as features. However, the long-range features of antigen proteins were not incorporated.

To process the long-distance information outside of peptides in antigenic proteins, several approaches considering amino acid sequences as series data were proposed. However, for example, an approach based on recurrent neural networks (RNN) [15] corresponding to a type of deep learning [16], did not include the amino acid sequence information outside an epitope. One of the related problems lied in inability to handle long-distance information appropriately, as it caused the problem of vanishing gradient during the learning process of RNN. In the present study, we consider the peptides themselves and the amino acid sequences before and after the peptides as the series data and address the problem of vanishing gradient by applying LSTM [6].

In addition, one of the essential problems associated with machine learning is that it is difficult to derive accurate predictions when the amount of training data is limited. In the previous study [5], physicochemical properties, such as type, composition, hydrophobicity, polarity, and the stability of amino acids in peptides, were employed as features in SVM. The proposed method further solves this problem by extending the model to include in a neural network the structural and chemical features of an entire antigen protein in addition to the amino acid sequence information.

## 4. Proposed method

### 4.1 Overview of proposed method

While predicting epitopes, the features and amino acid sequences within an epitope may not provide sufficient information for prediction. In the present study, we develop a method based on LSTM with an attention model in which the amino acid sequences inside and outside the epitope are considered as series data to handle the long-range information on proteins. In the following sections, we introduce LSTM and attention mechanism, and then describe the proposed method in detail.

### 4.2 LSTM

RNN [15] is a deep learning method that can be used to handle the series data. However, it is not applicable to the long-range series information due to problem of vanishing gradient, meaning that the gradient becomes small as a result of backpropagation during training.

LSTM [6] is an RNN model that can handle the long-range sequence information using memory cells and gates. Concerning memory cells, we can find that they pass only through simple addition and the Hadamard product by following a backpropagation flow. The backpropagation of simple addition is set to 1 on partial differentiation so that the gradient does not change, while the partial differentiation of the Hadamard product depends only on the output of a forgetting gate. That is, the gradient of an element that considered to be forgotten by the gate becomes small, while being maintained and transmitted in the past direction otherwise. Therefore, a memory cell can propagate the information that should be stored for a long time without losing it. In the present study, bi-directional LSTM [17] that is defined as a combination of the forward and backward LSTM models is implemented to combine the information obtained from the series corresponding to opposite directions.

### 4.3 Attention model

Although LSTM has the ability to retain the long-range information in contrast to RNN, the detailed information in an input series is tends to be lost, as it cannot be represented as a compressed vector. Therefore, we implement an attention mechanism that can refer directly to the information of an input series. Using such an attention mechanism, it is possible to memorize the vector output from an LSTM block at each series point and then multiply them by weights to obtain a context vector that defines what element in the context is to focus on.LSTM with an attention mechanism has achieved remarkable results in various application fields, including natural language processing [7]. In the present study, we introduce an attention mechanism to consider which parts of proteins should be emphasized in the model.

### 4.4 Utilization of the structural and chemical features of amino acid sequences

In general, the problem of machine learning methods is that they cannot be used to learn sufficiently in the cases when the amount of training data is limited. In the present study, we incorporate the structural and chemical features within peptides and whole antigenic proteins to enable the derivation of robust predictions even when the amount of data is sparse.

The structural and chemical features of the peptides considered in this study are the following: *β*-turn [18], relative surface accessibility [19], antigenicity [20], and hydrophilicity [21]. They are obtained using an epitope prediction application programming interface (API) provided by IEDB [9]. The total antigen protein features are considered as they are expected to affect the ease of binding between proteins and the epitopes. The following four features: isoelectric point, aromaticity, hydrophobicity, and stability, are obtained using the Biopython library [22]. A total of eight of the structural and chemical features are integrated into the network of the proposed method, as described the next section.

### 4.5 Epitope prediction of antigen protein using attention-based LSTM network

LSTM described in Section 4.2 is employed to preserve the long series information, and the attention mechanism introduced in Section 4.3 enables the prediction mechanism of notable locations. Then, incorporating the structural and chemical features, as suggested in in section 4.4 allows enabling robust estimation even when the amount of the training data is limited. In this section, we demonstrate how these three features can be combined in a single model to perform epitope predictions that are the subject of the present research.

In LSTM, the amino acid information at the point in a series and that before and after this point are combined into an embedded vector. It allows capturing the long-range amino acid sequence information.

Then, we apply the attention mechanism to estimate which amino acids are particularly noteworthy for the purposes of epitope estimation. For example, as the information corresponding to an inside peptide is considered to be more important than that associated with an outside one, it is possible to assign a higher weight to an inside peptide to propagate the information.

The final embedding vector obtained using LSTM with the attention mechanism and the structural and chemical features of the amino acid sequences, described in Section 4.4 are combined with the dense layer, and finally the presence of an epitope is judged using the sigmoid function. In the model learning phase, the model parameters at each learning step are employed to perform the above estimation, and then the model backpropagates the loss to epitope determination and updates the model parameters in each layer.

## 5. Evaluation experiments

### 5.1 Experimental conditions

To evaluate the applicability of the proposed methods, we compared with the baseline approach on the same dataset. In this experiment, we used BepiPred-2.0 [4] as the baseline model. Table 2 represents the comparison of the features considered in BepiPred-2.0 and in the proposed method.

**Table 2.** Comparison of the features used in BepiPred-2.0 and in the proposed method.

|  | BepiPred-2.0 | Proposed |
| --- | --- | --- |
| Embedding | Not used | Each amino acid in the target peptide and 16 amino acid before and after the peptide are embedded into a 20-dimensional vector. By concatenating the decoded vector (256-dimension) via LSTM and the attention decoder and peptide and protein features (8-dimension) as below, the output is obtained in the form of 264-dimensional embeddings. |
| Features about peptide | Molecular weight in amino acids, Hydrophobicity, Polarity | β-turn, Relative surface accessibility, Antigenicity, Hydrophilicity |
| Features about protein | Relative surface accessibility, Secondary structure | Isoelectric point, Aromaticity, Hydrophobicity, Stability |

Each model finally outputted a value in the range between 0.0 and 1.0 that indicates epitopicity. When this value was greater than 0.5, it is regarded as epitope, and when it was less than 0.5, it was considered as non-epitope. BepiPred-2.0 calculated predictions on the basis of amino acids, so that we used the average of the prediction scores for each amino acid as the predicted score of a target peptide. The trained model for BepiPred-2.0 was published as API [9], and we employed it to obtain result of evaluation, as described in Section 2.2.

We analyzed the performance concerning each antigen in terms of binary accuracy, similarly as [16]. To compare the performance of two models, the paired t-test was applied to their accuracy estimates on individual datasets. A confidence interval of 95% was considered to identify a significant difference between two compared models. Then, to focus on the prediction of positive labeling, we utilized the three indicators of positive labeling as described below:

- Sensitivity (Sens.) = True Positive / (True Positive + False Negative)
- Positive Prediction Value (PPV) = True Positive / (True Positive + False Positive)
- F1 value = 2 × PPV × Sensitivity / (PPV + Sensitivity)

Additionally, we evaluated the area under the curve (AUC) to adress the problem of bias in datasets. The chance rate in the present study was set as shown in the “Positive ratio” in Table 1 if all predictions were 1. Otherwise, it was set to 1− “positive ratio” if all predictions were 0. Notably, the chance rate was also high due to the bias in labels in the data.

### 5.2 Experimental results and discussions

The experimental results corresponding to the BepiPred-2.0 and the proposed methods are provided in Tables 3 and 4. As a reference, the results of the proposed method without the attention model (in each table, “w/o Attention”) were also included in a bottom line for each length.

**Table 3.** Accuracy and p-value of BepiPred-2.0 and the proposed methods. The highest scores among the compared methods in “Accuracy” columns and the value in “p-value” columns where there is a 5% significant difference are shown in bold. Macro average of accuracy for each method is shown in the first line as “Total.”

| Length | Method | Accuracy | p-value |
| --- | --- | --- | --- |
| Total | BepiPred-2.0 | 0.489 | — |
|  | proposed | **0.695** |  |
|  | w/o Attention | 0.673 |  |
| 8 | BepiPred-2.0 | 0.526 | 0.204 |
|  | proposed | 0.649 |  |
|  | w/o Attention | **0.774** |  |
| 9 | BepiPred-2.0 | 0.462 | **0.002** |
|  | proposed | **0.688** |  |
|  | w/o Attention | 0.638 |  |
| 10 | BepiPred-2.0 | 0.550 | **0.005** |
|  | proposed | **0.673** |  |
|  | w/o Attention | 0.657 |  |
| 11 | BepiPred-2.0 | 0.503 | **0.002** |
|  | proposed | **0.677** |  |
|  | w/o Attention | 0.644 |  |
| 12 | BepiPred-2.0 | 0.395 | **0.001** |
|  | proposed | **0.790** |  |
|  | w/o Attention | 0.582 |  |
| 13 | BepiPred-2.0 | 0.488 | **0.000** |
|  | proposed | **0.783** |  |
|  | w/o Attention | 0.708 |  |
| 14 | BepiPred-2.0 | 0.495 | **0.005** |
|  | proposed | **0.725** |  |
|  | w/o Attention | 0.705 |  |

**Table 4.** Sensitivity, positive prediction value, F1 value, and Area under the curve of the BepiPred-2.0 and proposed methods. The highest scores among the compared methods are shown in bold. Macro average of the score in each method is shown in the first line as “Total.”

| Length | Method | Sens. | PPV | F1 | AUC |
| --- | --- | --- | --- | --- | --- |
| Total | BepiPred-2.0 | 0.384 | **0.768** | 0.479 | 0.569 |
|  | proposed | **0.775** | 0.725 | **0.705** | 0.706 |
|  | w/o Attention | 0.644 | 0.752 | 0.656 | **0.713** |
| 8 | BepiPred-2.0 | 0.530 | **0.736** | 0.546 | 0.576 |
|  | proposed | **0.858** | 0.547 | 0.610 | **0.822** |
|  | w/o Attention | 0.733 | 0.640 | **0.644** | 0.801 |
| 9 | BepiPred-2.0 | 0.337 | **0.898** | 0.479 | 0.595 |
|  | proposed | **0.819** | 0.788 | **0.790** | 0.626 |
|  | w/o Attention | 0.697 | 0.839 | 0.732 | **0.709** |
| 10 | BepiPred-2.0 | 0.285 | 0.449 | 0.311 | 0.499 |
|  | proposed | **0.393** | 0.578 | **0.400** | **0.792** |
|  | w/o Attention | 0.292 | **0.580** | 0.350 | 0.745 |
| 11 | BepiPred-2.0 | 0.409 | **0.980** | 0.574 | **0.694** |
|  | proposed | **0.763** | 0.838 | **0.780** | 0.628 |
|  | w/o Attention | 0.682 | 0.855 | 0.744 | 0.693 |
| 12 | BepiPred-2.0 | 0.339 | 0.689 | 0.409 | 0.470 |
|  | proposed | **0.698** | **0.790** | **0.676** | **0.739** |
|  | w/o Attention | 0.557 | 0.736 | 0.581 | 0.619 |
| 13 | BepiPred-2.0 | 0.406 | **0.900** | 0.553 | 0.605 |
|  | proposed | **0.951** | 0.805 | **0.871** | 0.661 |
|  | w/o Attention | 0.752 | 0.859 | 0.777 | **0.716** |
| 14 | BepiPred-2.0 | 0.381 | 0.726 | 0.484 | 0.545 |
|  | proposed | **0.940** | 0.730 | **0.806** | 0.677 |
|  | w/o Attention | 0.796 | **0.756** | 0.764 | **0.712** |

**Table 5.**
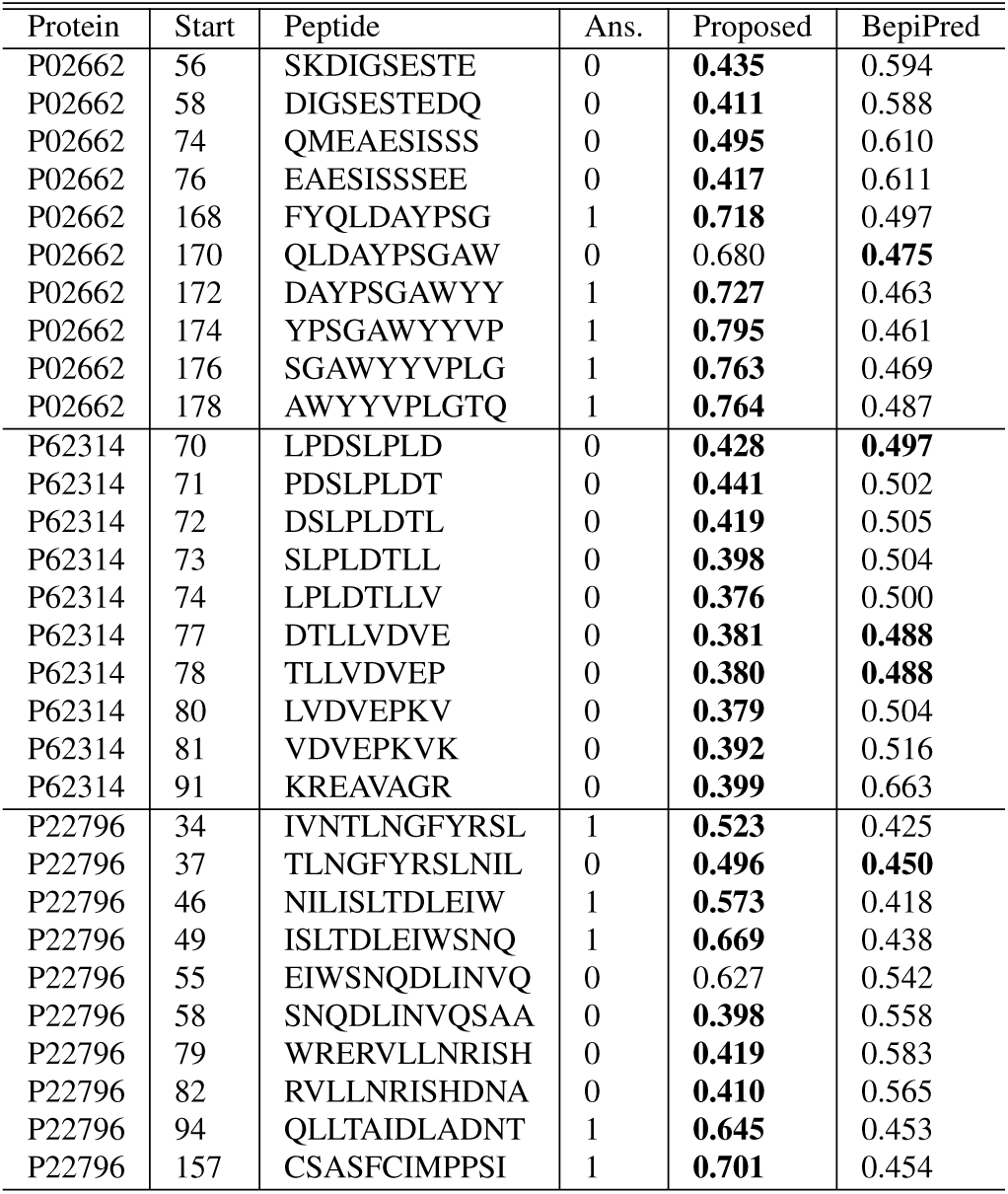
Example of epitope prediction: “Start” denotes the start position of peptide. “Ans.” is an answer label where 1 indicate that target peptide is epitope. Each value in the “Proposed” and “BepiPred-2.0” columns represents an estimated value, and estimated value is marked in bold if it matched to an answer label.

We observed that the proposed method achieved the highest accuracy concerning all lengths compared to BepiPred-2.0, as represented in Table 3.

These results indicated that the proposed method significantly outperformed BepiPred-2.0 except for length 8, when p-values below 0.05 were considered statistically significant. Moreover, as shown in Table 4, the proposed method outperformed BepiPred-2.0 in terms of sensitivities, F1 scores, and AUC for almost all lengths. Although the scores at particular lengths were below BepiPred-2.0 in terms of PPV, the AUC scores of the proposed model were higher than those of BepiPred-2.0 in almost all length. Therefore, we concluded that there was room for improvement through adjusting the threshold.

Next, we compared the results of the proposed method with and without the attention model. Although only a small difference in PPV was observed between them, the sensitivity was less than that of the proposed method, suggesting that the importance of the attention model could play an important role in epitope prediction. These issues are planned to be addressed in the future research.

### 5.3 Example of detection by the proposed method

In this section, we discuss a prediction case for the proposed method. First, we considered the predicted results of BepiPred-2.0 and LSTM, including the peptides with no label data existing, as represented in Figures 1 and 2, respectively. As an example of a target protein, a 10-length peptide in protein P02662 was analyzed. The horizontal axis in these figures represents the starting point of a peptide in a protein, and the vertical axis denotes the predicted value of the target peptide. Here the, blue, and gray maps indicate the positive datay, negative data, and no data for a label respectively. For example, a plotting point at the point, where the horizontal axis is 1 denotes the point where the amino acids from the 1st one to 10th are regarded as a peptide representing an estimation value, and it does not has the label data in dataset.

**Fig. 1.**
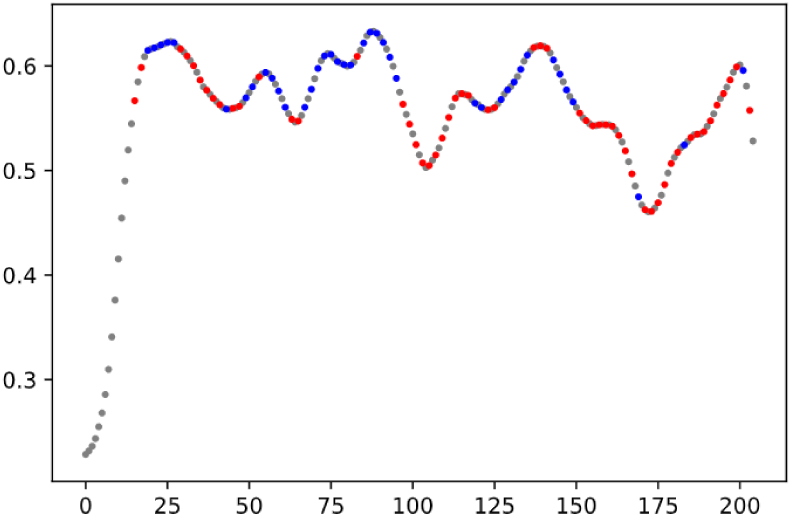
Plot of the predicted value of protein P02662 by BepiPred-2.0.

**Fig. 2.**
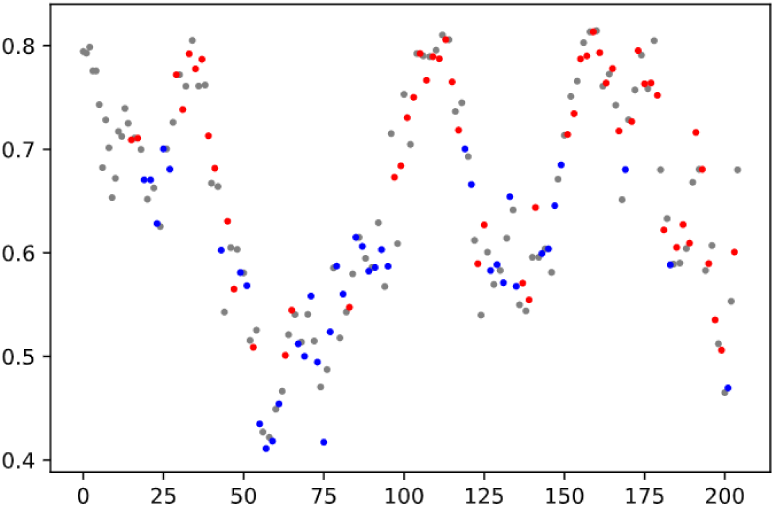
Plot of the predicted value of protein P02662 by the proposed method.

The results of BepiPred-2.0, as represented in Figure 1 demonstrated that the positive red and negative blue examples tended to be placed in opposite positions for corresponding to the movements of the peaks and valleys of the predictions on the vertical axis. However, concerning the proposed method, as shown in Figure 2, it could be seen that the tendency of peaks and valleys, and the tendency of the red/blue color classification were relatively similar. In addition, if the threshold is determined appropriately, rather than being set identical (0.5) for all proteins, it is expected that the accuracy may be improved further.

Table 4 represents the peptides and their correct and estimated values concerning the three protein examples: P02662, P62314, and P22796. Here, P02662 is the same target as Figures 1 and 2. As indicated in Table 4, the proposed method was able to recognize a peptide as non-epitope, unlike BepiPred-2.0. Concerning P62314, the answer labels were often negative; however BepiPred-2.0 incorrectly gave a high estimated score, while the proposed method outputted the negative one. However, concerning the peptides QLDAYPSGAW in P02662 and EIWSNQDLINVQ in P22796, the proposed method also provided negative incorrectly. It is necessary to investigate and mitigate the causes of such cases of incorrect estimation in the future.

## 6. Conclusion

In the present paper, we proposed a new model for B-cell epitope prediction considering the following features:

1. Not only peptides but also proteins were processed as the amino acid sequence data and were modeled using an LSTM method based on attention model.
2. The combination of the structural and chemical features of peptides and whole proteins enabled robust predictions even on a limited amount of learning data.

We demonstrated that the proposed model achieved superior performance compared with the existing method (BepiPred-2.0). This proposed method can also be applied to the prediction of interactions between the partial and whole protein sequences.

The issues to be addressed in the future research include the need to confirm of the effectiveness of the proposed method through conducting experiments based on applying it to antibody proteins other than IgG, as well as through considering the case when the structural and chemical features of an amino acid sequence are unified with the comparative method. In the experiments, we excluded peptides which were either too short or too long from the experiment for stable training. Dealing with the prediction of peptide with chain lengths other than 8-14 will be considered as an issue to focus on in the future research.

Furthermore, we will focus on modeling the protein three-dimensional structures used for the prediction of nonlinear epitopes [25] that are deemed to be a predicting source of information. In addition, we would like to further improve the prediction accuracy of the proposed method by utilizing the protein three-dimensional structures. Dataset and some basic programs will be made available to the public.

## Footnotes

*1 https://www.kaggle.com/futurecorporation/epitope-prediction

